# CRISPR-Tag: an Efficient DNA Tagging System in Living Cells

**DOI:** 10.1101/280495

**Authors:** Baohui Chen, Wei Zou, Bo Huang

## Abstract

A lack of efficient tools to image non-repetitive genes in living cells has limited our ability to explore the functional impact of spatiotemporal dynamics of genes. Here, we addressed this issue by developing the CRISPR-Tag system as a new DNA tagging strategy to label protein-coding genes with high signal-to-noise ratio under wild-field fluorescence microscopy by using 1 to 4 highly active sgRNAs. The CRISPR-Tag, with minimal size of ∼ 250 bp, represents an easily and broadly applicable technique to study spatiotemporal organization of genomic elements in living cells.

---

The CRISPR-Cas9 system has been reprogrammed to label specific genomic loci in living cells. When targeting non-repetitive genomic regions, it requires multiple sgRNAs function simultaneously to provide sufficient signal-to-noise ratio for microscopy detection^1^. Although the number of sgRNAs could be reduced to 3 to 4 using a combination of signal amplification and super-resolution microscopy^2^,^3^, the labeling efficacy has not been quantitatively assessed. It is also important to note that the efficiency of Cas9 targeting for any genomic locus can be dramatically influenced by the activity of sgRNAs used^4^. As such, it is very likely that only part of sgRNAs selected for DNA labeling function with high activity, which remains the major uncertainty of CRISPR-mediated genomic labeling. Thus, well-designed approaches using CRISPR imaging as readouts are critical to further optimize the DNA labeling system. Collectively, the major challenge is to develop an efficient DNA tagging system to achieve the full potential of CRISPR imaging technology for labeling non-repetitive genomic elements.

To overcome this limitation, we developed a CRISPR-DNA tag system, termed “CRISPR-Tag”, to achieve efficient labeling of non-repetititve genes (diagram in **Fig. 1a**). CRISPR-Tag was assembled with CRISPR targeting sequences from *C. elegans* genome which have been characterized for genome editing by several studies^5^-^8^. Six target sequences were picked according to the editing efficiency in worms and the on/off-target activity prediction by the web tool (http://crispr.mit.edu/). In addition, we generated a piece of artificial sequence based on the preference of nucleotides sequences that impact sgRNA efficacy^9^. In total, seven sgRNAs, termed sgTS1 to sgTS7, were selected as the candidate sequences to assemble CRISPR-Tags (**Table 1**). The first version of CRISPR-Tag (CRISPR-Tag_v1) contains six repeats. Four CRISPR targeting sequences, TS1 to TS4 were arranged in each repeat unit. To label a specific non-repetitive gene, we aim to first insert CRISPR-Tag into its 3’ UTR region or intron region by CRISPR knock-in, and then label the CRISPR-Tag with the nuclease-deficient Cas9 (dCas9) fused with fluorescent tags (**Fig. 1a**).

**Figure 1.**
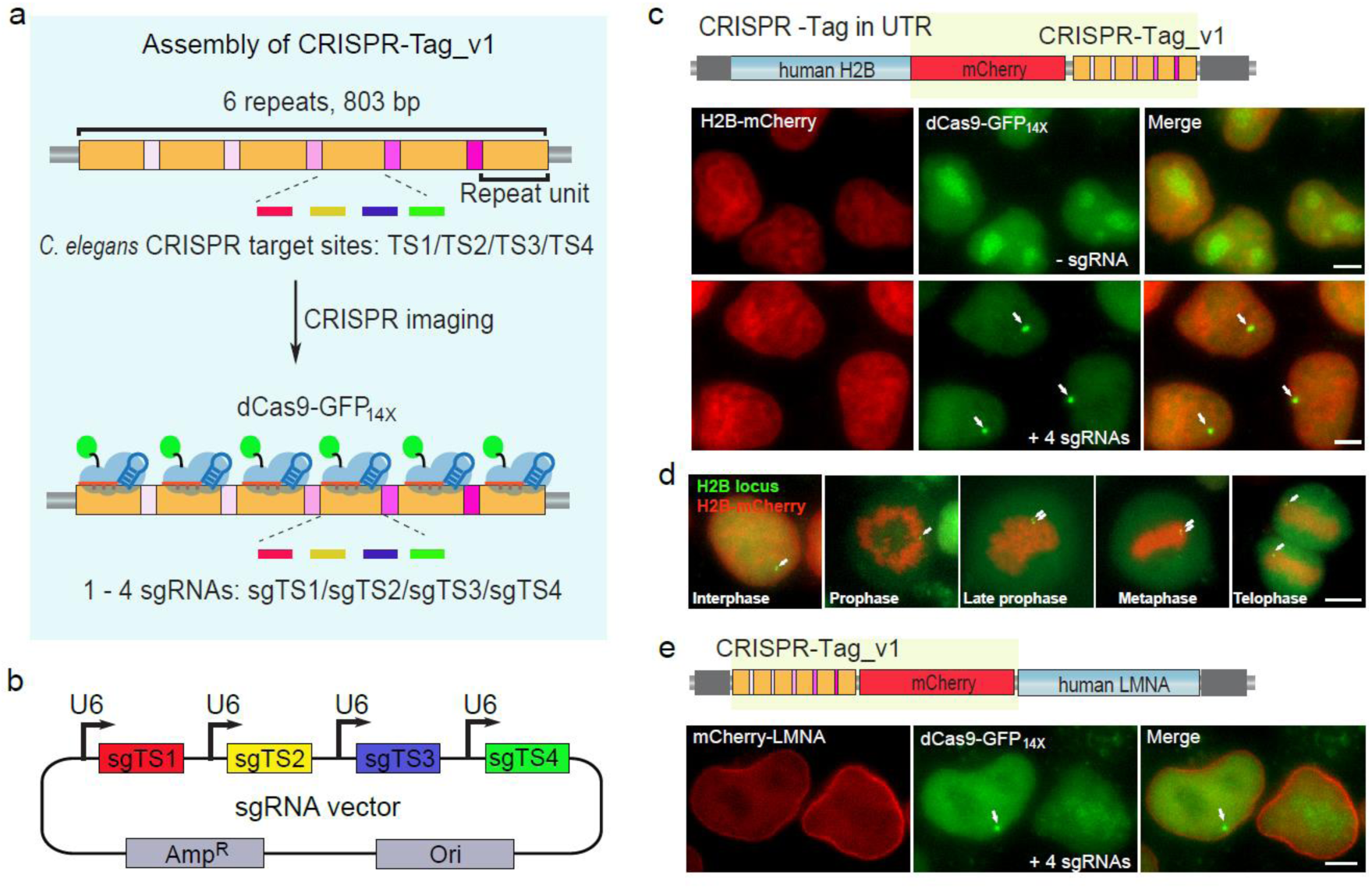
Development of CRISPR-Tag to label non-repetitive genes. (a) Schematic of CRISPR-Tag design as a new DNA tagging system. (b) Co-expression of four sgRNAs in one vector. sgTS1, sgTS2, sgTS3 and sgTS4 were built individually and then sub-cloned into a single vector. (c) mCherry-CRISPR-Tag_v1 was inserted into C-terminus of human H2B gene as highlighted by yellow. Representative images denoted simultaneous visualization of H2B-mCherry and H2B loci by using CRISPR-Tag. H2B loci were labeled by dCas9-GFP_14X_ with 4 sgRNAs (sgTS1 to sgTS4). (d) H2B-mCherry and H2B loci were visualized at different stages of cell cycles. (e) Representative images to show simultaneous visualization of mCherry-LMNA and LMNA loci by utilizing the CRISPR-Tag system. All images are maximum intensity projections from z stacks. Scale bars: 5 μm.

In order to minimize the size of the CRISPR-Tag, i.e. the number of target sequences, required to generate detectable signal, we adopted the tandem split GFP system to amplify the fluorescence from dCas9^10^. To this end, we fused a repeating array containing 14 copies of GFP11 tags to dCas9 (dCas9-GFP_14x_), which can theoretically recruit as many as 14 copies of GFP (**Supplementary Fig. 1a**). When labeling the repeats in *MUC4* and 5S rDNA genes and imaging with wild-field fluorescence microscopy, dCas9-GFP_14x_ increased the signal-to-noise ratio (SNR) by a factor of 3 compared to dCas9-EGFP (**Supplementary Fig. 1b,c**), demonstrating an enhanced microscopy detection efficiency. However, when imaging non-repetitive regions in *MUC4* gene with our previously published 36 sgRNAs that have not been individually validated, signal amplification alone by dCas9-GFP_14x_ still gives relatively low SNR (**Supplementary Fig. 2**). This result further indicates the need for a small set of well validated sgRNAs as in our CRIPSR-Tag system.

**Figure 2.**
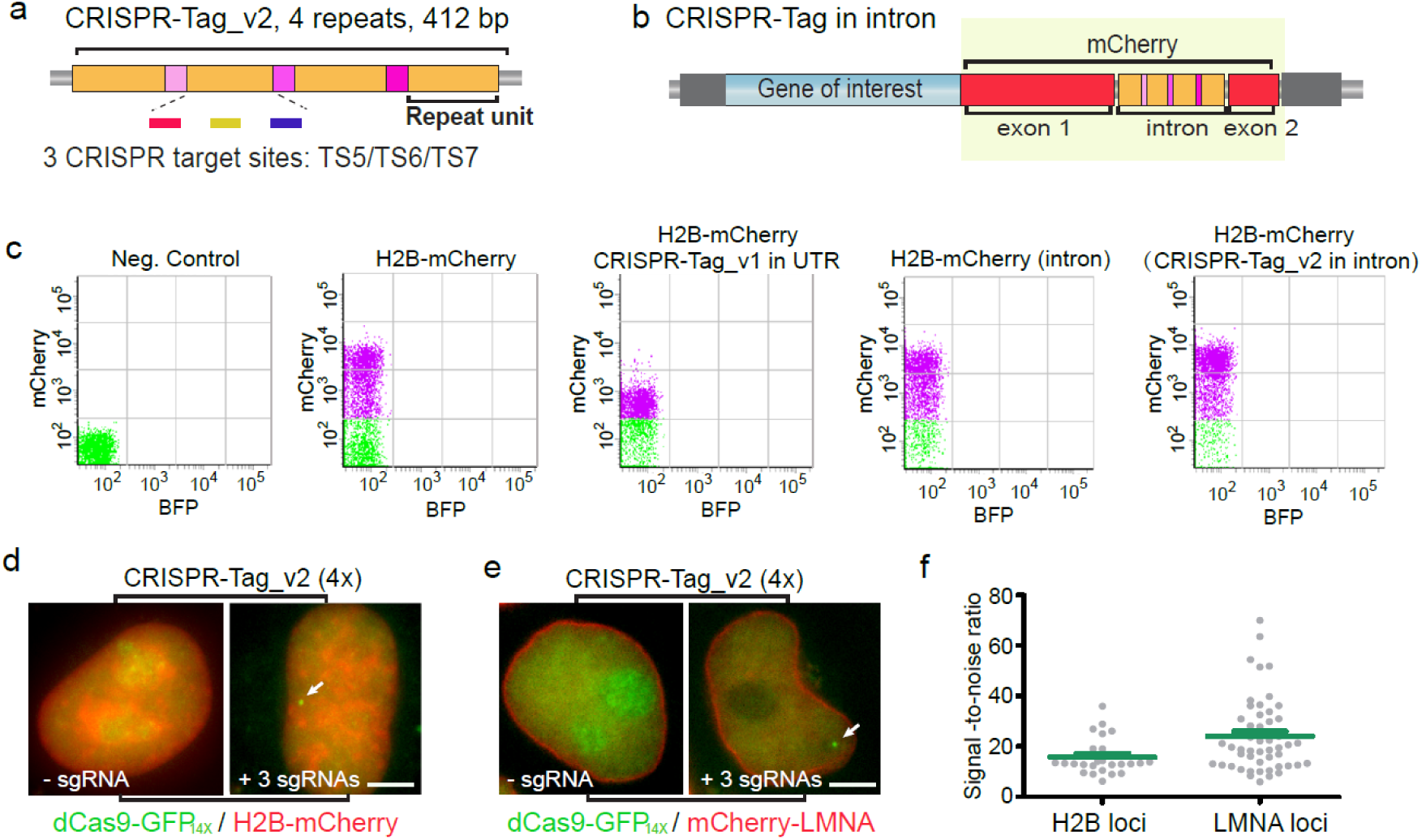
The protein expression remains normal when hiding CRISPR-Tag in intron. (a) Schematic of CRISPR-Tag_v2 design which contains 4 repeats. Each repeat harbors 3 CRISPR targeting sites including TS5, TS6 and TS7. (b) Schematic of CRISPR-Tag positioning in the intron of mCherry. The fragment including mCherry and CRISPR-Tag, indicated by yellow, was integrated to the N- or C-terminus of a target gene. (c) FACS analysis was performed to characterize H2B-mCherry expression in HeLa cells. Each dot represents single cells. BFP serves as an irrelevant channel. The distribution of dots along the y axis indicates mCherry signal intensity. (d) Visualization of H2B locus using CRISPR-Tag_v2 which was inserted into the intron of mCherry. H2B-mCherry was imaged and H2B loci were labeled by dCas9-GFP_14X_ with 3 sgRNAs (sgTS5 to sgTS7). (d) Visualization of LMNA locus using CRISPR-Tag_v2 with 3 sgRNAs (sgTS5 to sgTS7). (e) Labeling efficiency of H2B and LMNA loci shown in (d) and (e), respectively, was determined by quantifying signal-to-noise ratio, n ≥ 25 cells. Green line denotes means ± SEM.

To demonstrate our CRISPR-Tag, we chose a protein gene as our test system in order to simplify the selection process of CRISPR-Tag knock-in cells. Specifically, we inserted CRISPR-Tag into the 3’ UTR region of human H2B by CRISPR knock-in using electroporation of Cas9/sgRNA ribonucleoprotein complexes (RNPs)^11^ and double-cut plasmid donor^12^, which could increase targeting specificity and efficiency, respectively (diagram in **Supplementary Fig. 3**). We found that double-cut plasmid donor indeed resulted in higher knockin efficiency than conventional plasmid donor and the efficiency highly dependents on cell types (**Supplementary Fig. 4a,b**). To select the integrated cells, we further added a mCherry sequence to the tag, which was knocked into the C-terminus of H2B as a FACS sorting marker (**Supplementary Fig. 4c,d**). CRISPR-Tag insertion was validated by PCR (**Supplementary Fig. 5**) and was further confirmed by the subcellular localization of H2B-mCherry. We then performed CRISPR imaging experiments using dCas9-GFP_14x_ Tag. As expected, one bright puncta, representing H2B locus, was clearly visible in most cells upon transfection of four sgRNAs (sgTS1-sgTS4) (**Fig. 1b,c**). Histone H2B, as one of the 5 main histone proteins, is involved in the formation of chromatin structure in eukaryotic cells^13^. H2B-mCherry allowed imaging of both interphase chromatin and mitotic chromosomes and revealed various chromatin states. In the meanwhile, we observed that H2B locus underwent replication and separation into two daughter cells at the telophase stage (**Fig. 1d**). Similarly, we achieved simultaneous imaging of protein and gene positioning for both human nuclear membrane protein LMNA, and heat-shock response protein HSPA8 in living cells (**Fig. 1e; Supplementary Figs. 4 and 6**).

**Figure 3.**
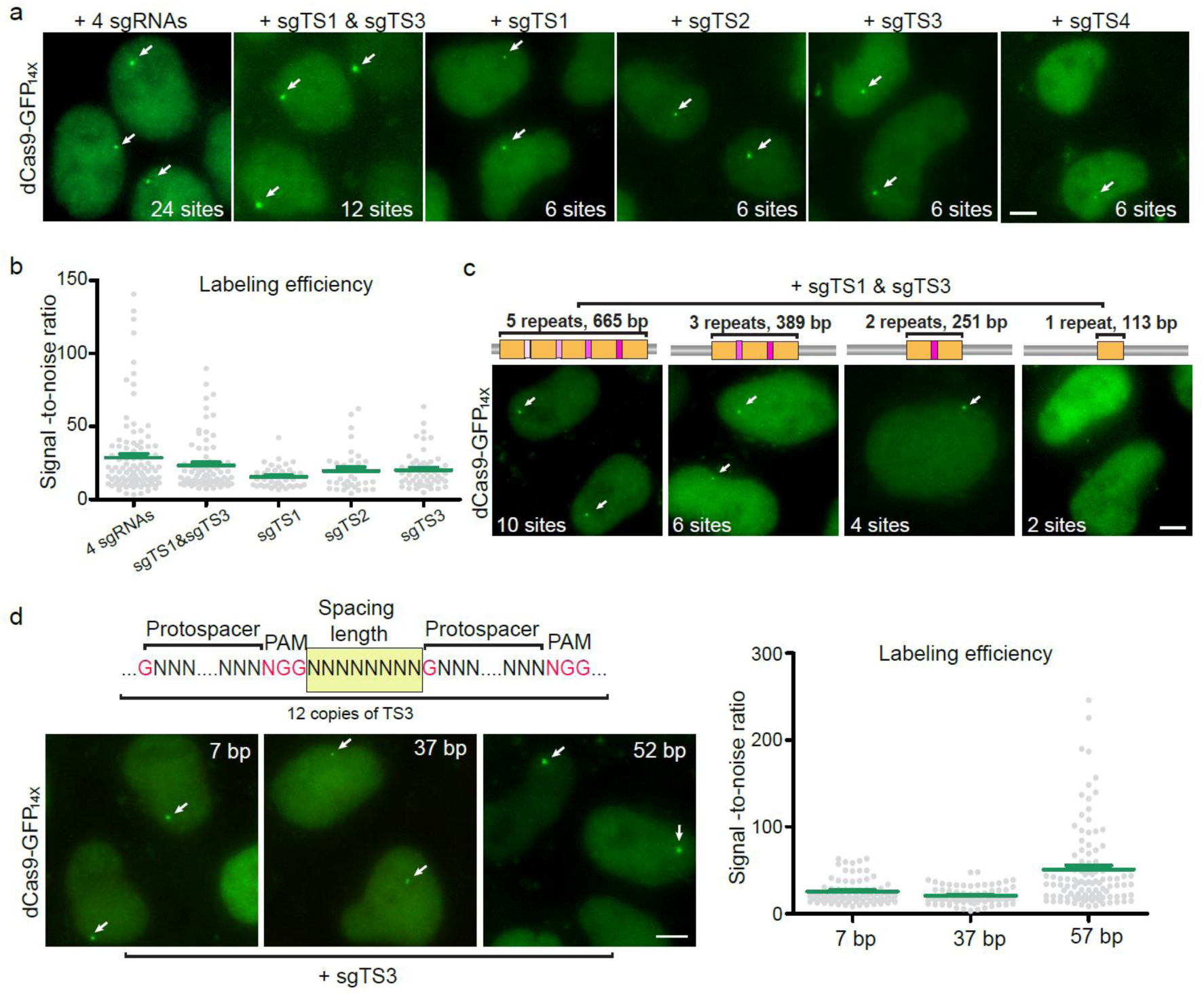
Assess CRISPR-Tag designs for effective DNA labeling. (a) Minimal number of sgRNAs to label H2B loci with CRISPR-Tag_v1. The use of 4 sgRNAs allows 24 binding sites of dCas9-GFP_14X_ and one sgRNA facilitates 6 CRISPR targeting sites. Representative images were shown for each case. Detected H2B loci are indicated by arrows. (b) Comparison of labeling efficiencies of using different sgRNAs, n ≥ 33 cells. Error bars report ±SEM. (c) Minimal number of CRISPR targeting sites to label H2B loci. CRISPR-Tag containing different copies of repeat units were generated and then labeled by expressing both sgTS1 and sgTS3, respectively. (d) Optimal spacing length between two neighboring targeting sites. Spacing length was defined as the number of nucleotides from NGG to next protospacer sequence as indicated in the diagram. Labeling efficiency was defined by quantifying signal to noise ratio, n ≥ 72 cells. Green line reports means ± SEM. All images in Fig. 2 are maximum intensity projections from z stacks. Scale bars: 5 μm.

Specifically when tagging protein-coding genes, it is possible that certain 3’ UTR sequences may affect the expression level. Indeed, comparing to inserting mCherry alone, inserting mCherry+CRISPR-Tag caused a decrease in protein expression level for H2B, LMNA and HSPA8 in HeLa cells (**Supplementary Fig. 7a-c**) but not H2B in 293T cells (**Supplementary Fig. 7d**). To minimalize this effect, we engineered the mCherry coding sequence to add in an intron while maintaining its expression level (**Supplementary Fig. 8a,b**). We have also assembled a second version of CRISPR-Tag, termed CRISPR-Tag_v2, which can be recognized by sgTS5, sgTS6 and sgTS7 (**Fig. 2a,b)**. When inserting in the mCherry intron, CRISPR-Tag_v2 had no effect on H2B-mCherry expression, whereas our earlier design CRISPR-Tag_v1 caused a decrease in mCherry expression, possibly due to its interference with spicing (**Fig. 2c; Supplementary Figs. 8c,d and 9**). Then, we validated this new tag for the labeling and imaging of H2B and LMNA genes. With this new strategy, both H2B and LMNA genes could be efficiently visualized using three sgRNAs while not affecting their expression levels (**Fig. 2d-f**). Our results showed that CRISPR-Tag_v2 could be as short as 412 bp containing four repeats. We then measured the positioning of *H2B* and *LMNA* loci in the nucleus and found both *H2B* and *LMNA* genes preferentially locate closer to the nuclear periphery than to the nuclear center (**Supplementary Fig. 10**). These results reveal the nonrandom spatial organization of allele-specific locus in the nucleus.

We next performed a series of optimizations and quantifications in order to reduce the size of the CRISPR-Tag while maintaining its labeling efficiency. First, we compared signal-to-noise ratio by using different numbers of sgRNAs to label CRISPR-Tag_v1, which is 803 bp long containing six repeats. Each repeat can be recognized by up to four sgRNAs (sgTS1 to sgTS4). We observed that transfection of one single sgRNA, such as sgTS1, sgTS2 and sgTS3, resulted in sufficiently visualization of CRISPR-Tag with SNR above 15. The SNR increased to 23.3 ± 8.7 and 28.6 ± 10.1 (mean ± SEM) when using two and four sgRNAs respectively (**Fig. 2a,b**). Collectively, one sgRNA to target six sites is sufficient for labeling a specific locus. We next sought to further reduce the size of CRISPR-Tag by assembling fewer repeats, including 5, 3, 2 and 1 repeats. We found that the size of CRISPR-Tag can be as small as ∼250 bp harboring 4 CRISPR targeting sites (**Fig. 2c**). It is also important to optimize the spacing of two adjacent targeting sites in the CRISPR-Tag because steric hindrance of neighboring dCas9 protein binding sites is a concern. We varied the spacing length and quantified SNR. The results indicated that 52 bp spacing linker separating the two neighboring targeting sites achieved the best result (**Fig. 2d**). Together, our studies provide some critical guidelines for CRISPR-Tag design. The smallest CRISPR-Tag is substantially shorter than the LacO/TetO array (usually ∼10 and ∼4 kb, respectively)^14^, and the ParB/INT (∼ 1 kb) system^15^. Important to note, CRISPR-Tag is much more versatile than these DNA tags, as a wide range of orthogonal CRISPR-Tags with different CRISPR targeting sequences can be assembled.

In conclusion, by assembling *C. elegans* genomic sequences which can be effectively targeted by the CRISPR-Cas9 system, we created CRISPR-Tag as a new DNA tagging tool in living human cells. CRISPR-Tag, as small as 250 bp, enables live-cell labeling of non-repetitive genes at the level of single cells and single loci. In theory, this technique can be applied broadly for any other species, such as using Human CRISPR-Tag for *C. elegans* genomic loci labeling. Furthermore, a same set of CRISPR-Tag and sgRNAs is applicable for tagging any genomic loci. In addition, CRISPR-Tag can be also designed as a non-repetitive DNA tag or assembled with other unique sequences for new tool developments. As a proof of concept, we have showed that CRISPR-Tag enables efficient labeling of protein-coding genes. However, this new technique can be potentially adapted for labeling regulatory elements in the genome. Thus, CRISPR-Tag opens up possibilities for both new genomic tool developments and biological applications.

## Material and methods

### Cell culture

HeLa and HEK293T cells were maintained in Dulbecco’s modified Eagle medium (DMEM) with high glucose (Gibco) in 10% FBS (Clontech) and 1% penicillin/streptomycin (Gibco). All cells were grown at 37 °C and 5% CO_2_ in a humidified incubator.

### Construction of dCas9 and sgRNA

The nuclease-deficient *Streptocococcus pyogenes* Cas9 (dCas9) was used for all the CRISPR imaging experiments in this study. The construction of dSpCas9-EGFP has been previously described ^1^. To build dCas9-GFP11_14x_, we first synthesized two fragments of gBlocks, each containing 7 copies of the coding sequence of GFP11 tag. Then we modified the SunTag vector (pHR-dSV40-NLS-dCas9-24xGCN4_v4-NLS-P2A-BFP-dWPRE, addgene #60910) to replace 24xGCN4 with GFP11_14x_. To express GFP1-10 in the nucleus, we fused GFP1-10 with one copy of SV40 NLS and cloned the fusion protein into a lentiviral vector containing a strong promoter *P*_SFFV._ Construction of sgRNAs to label MUC4 and sg5S rDNA genes (**Supplementary Fig. 1 and Fig. 3**) in a lentiviral vector has been described in our previous study (1-2). Other sgRNAs in this study, including sgTS1 to sgTS7, were built by modifying the CRISPRainbow vector (addgene #75398). For co-expression of multiple sgRNAs (>2), the Golden Gate Cloning method was used to assemble multiple sgRNAs into a same vector as described previously^16^.

### Assembly of CRISPR-Tag

To assemble the CRISPR-Tag, a gBlock containing a series of *C. elegans* genomic sequences (GNNNNNNNNNNNNNNNNNNN<u>NGG</u>) that can be recognized by CRISPR-Cas9 system was synthesized by Integrated DNA Technologies (IDT). The spacing of two neighboring CRISPR targeting sits varies according to the design in Fig. 1 and Fig. 2. To assemble a CRISPR-Tag with a desired number of repeats, the repeat unit containing 3 or 4 CRISPR targeting sites was amplified by PCR reactions using the Phusion High-Fidelity DNA polymerase (New England Biolabs). The Golden Gate Cloning method was then performed to assemble CRISPR-Tags that consist of different desired numbers of repeats. Important to note, there is a unique sequence arranged between two adjacent repeats in the CRISPR-Tag, which facilitates easy validation of CRISPR knock-in by PCR reactions.

### Re-engineering mCherry for carrying the CRISPR-Tag

The fourth intron of human HSPA5 gene was inserted into mCherry-coding sequence right after the three nucleotides that code for the 197^th^ amino acid of mCherry protein. A restriction site BstXI was artificially embedded in the intron region for the ease of molecular cloning. CRISPR-Tag was then cloned into the BstXI site by In-Fusion HD Cloning Kit (Clontech).

### Construction of donor plasmids

All donor plasmids used in this study were generated with NEBuilder HiFi DNA Assembly Cloning Kit (New England Biolabs). For example, to construct the donor plasmid for inserting mCherry and CRISPR tag to the C-terminus of H2B, the left and right homology arm (HA) were amplified from the genomic DNA of HeLa cells, with the stop codon being removed. Other two fragments, mCherry and CRISPR tag were amplified respectively. Then the four fragments, including left HA, mCherry, CRISPR-Tag and right HA were assembled into a same vector. To generate donor plasmids harboring sgH2B recognition sites, termed double-cut donor plasmid, the sgH2B targeting sequence together with the PAM sequence (GCGAGCGCCAGGTCCCGGCA<u>GGG</u>) was included in the forward primer of left HA and the reverse primer of right HA. Therefore, sgH2B targeting sequence was tagged to the regions flanking the upstream and downstream HA as previously described^12^.

### Lentiviral production

HEK293T cells were seeded into 6-well plates 24 hours prior to transfection. 110 ng of pMD2.G plasmid, 890 ng of pCMV-dR8.91 and 1000 ng of the lentiviral vector (dCas9-EGFP, dCas9-GFP11_14x_, GFP1-10 or sgRNA) were co-transfected into HEK293T in each well using FuGENE (Promega) following the manufacture’s recommended protocol. Virus was harvested 48 hours after transfection.

### Transfection, infection and generation of stable cell lines

HeLa cells were infected with dCas9-EGFP and Tet-on 3G lentiviruses, and then clonal cell lines were isolated to expressed dCas9-EGFP at a suitable level. A more detailed protocol to isolate clonal cell lines of dCas9-FP has been described^17^.The basal expression level of dCas9-EGFP without doxycycline induction is ideal to achieve high signal-to-noise ratio. To generate HeLa cells stably expressing dCas9-GFP_14x_, we infected HeLa cells with dCas9-GFP_14x_ and GFP1-10 lentivirues. A clonal cell line which achieved the best signal-to-noise ratio was selected for CRISPR imaging. The non-repetitive region of *MUC4* gene was labeled by infecting dCas9-FP expressing cells with a mixed lentiviral cocktails containing 26 sgRNAs. To label repetitive genomic elements or specific protein-coding genes, dCas9-FP stable cell line without /with successful knock-in of CRISPR-Tag was transfected with sgRNAs using FuGENE (Promega) following the manufacture’s recommended protocol. For all the CRISPR imaging experiments, HeLa cells were plated into 8-well chambered coverglass (Lab-Tek II). 400 ng of total sgRNA plasmid DNA were transfected for each individual well.

### sgRNA in vitro transcription

sgRNAs for CRISPR-mediated knock-in was transcribed in vitro following the published protocol^18^. The following sequence was used as the DNA template to transcribe sgRNAs in vitro: 5’-TAA TAC GAC TCA CTA TAG GNN NNN NNN NNN NNN NNN NNG TTT AAG AGC TAT GCT GGA AAC AGC ATA GCA AGT TTA AAT AAG GCT AGT CCG TTA TCA ACT TGA AAA AGT GGC ACC GAG TCG GTG CTT TTT TT-3’ containing a T7 promoter (TAATACGACTCACTATAG), a gene specific ∼20-nt protospacer sequence starting with a G for optimal T7 transcription (GNNNNNNNNNNNNNNNNNNN), and a common sgRNA scaffold region. The DNA template was generated by overlapping PCR using a set of four primers: three primers common to all reactions (forward primer T25: 5’-TAA TAC GAC TCA CTA TAG-3’; reverse primer BS7: 5’-AAA AAA AGC ACC GAC TCG GTG C-3’ and reverse primer ML611: 5’-AAA AAA AGC ACC GAC TCG GTG CCA CTT TTT CAA GTT GAT AAC GGA CTA GCC TTA TTT AAA CTT GCT ATG CTG TTT CCA GCA TAG CTC TTA AAC-3’) and one gene-specific primer (forward primer 5’-TAA TAC GAC TCA CTA TAG GNN NNN NNN NNNNNN NNN NNG TTT AAG AGC TAT GCT GGA A-3’). For each template, a 100 μL PCR was set using iProof High-Fidelity Master Mix (Bio-Rad) reagents. The PCR product was then purified using DNA Clean and Concentrator-5 columns (Zymo Research) following the manufacturer’s instructions. Next, a 100 μL in vitro transcription reaction was carried out using HiScribe T7 High Yield RNA Synthesis Kit (New England Biolabs). The sgRNA product was then purified on RNA Clean and Concentrator-5 columns (Zymo Research) and eluted in 15 μL of RNase-free RNA buffer (10 mM Tris pH 7.0 in DEPC-treated H2O). sgRNA quality was examined by running 3 pg of the purified sgRNA on a 10% polyacrylamide gel containing 7 M urea (Novex TBE-urea gels, ThermoFisher Scientific).

### CRISPR-mediated knock-in

RNP assembly and electroporation were performed to carry out CRISPR knock-in experiments. Cas9/sgRNA RNP complexes were prepared following methods reported by Lin et al.^11^. Cas9 protein (pMJ915 construct, containing two nuclear localization sequences) was expressed in E. coli and purified by the University of California Berkeley Macrolab following protocols described by Jinek et al.^19^. The HeLa dCas9-GFP_14x_ cells were treated with 200 ng/mL nocodazole (Sigma) for 15 h prior to electroporation to increase HDR efficiency as shown by Lin et al. (14). RNP complexes were assembled with 100 pmol Cas9 protein and 130 pmol sgRNA just before electroporation and combined with 400 ng donor plasmid DNA in a final volume of 10 μL. First, 130 pmol purified sgRNA was diluted in Cas9 buffer (final concentrations: 150 mM KCl, 20 mM Tris pH 7.5, 1 mM TCEP-HCl, 1 mM MgCl2, 10% vol/vol glycerol) and incubated at 70 °C for 5 min. A total of 2.5 μL of Cas9 protein (40 μM stock in Cas9 buffer, i.e., 100 pmol) was then added and RNP assembly was carried out at 37 °C for 10 min. Finally, donor plasmid DNA was added to this RNP solution. Electroporation was performed in a Amaxa 96-well shuttle Nucleofector device (Lonza) using SF-cell line reagents (Lonza) following the manufacturer’s instructions. Nocodazole-treated HeLa cells were washed with PBS and resuspended to 10^4^ cells per microliter in SF solution immediately before electroporation. For each sample, 20 μL of cells (i.e., 2 ×10^5^ cells) was added to the 10 μL RNP/template mixture. Cells were immediately electroporated using the CM130 program and transferred to 1 mL pre-warmed culture medium in a 24-well plate. FACS selection of knock-in positive cells was carried out 3 days after electroporation.

### Flow cytometry

3 days following Cas9/sgRNA/donor electroporation, cells were analyzed by flow cytometry on aLSR II instrument (BD Biosciences) and sorted on a FACS Aria II (BD Biosciences). Cells were first gated for the intact cell population using forward scatter versus side scatter plots and then gated for single cells based on forward scatter W versus forward scatter H. mCherry positive cells were sorted out for further validation of CRISPR knock-in.

### Microscopy and data analysis

CRISPR imaging data were acquired on a Nikon Ti-E inverted wide-field fluorescence microscope equipped with a 100x NA 1.40 PlanApo oil immersion objective, an LED light source (Excelitas X-Cite XLED1), an sCMOS camera (Hamamatsu Flash 4.0), and an motorized stage (ASI) with stage incubator (Tokai Hit). Acquisitions were controlled by MicroManager. All images were taken as z stacks at 0.4 μm steps and with a total of 15 steps and were projected in maximum intensity. All the fluorescence imaging data were analyzed by Image J. Signal-to-noise ratio was defined as the ratio of the intensity of a fluorescent signal and the power of background noise as following formula:

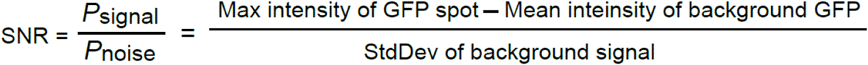

## Acknowledgements

We thank D. Kamiyama, S. Sekine, M. Leonetti, S. Feng and Y. Wang for fruitful discussions. The studies in Chen lab are supported by The Thousand Talents Plan Startup. The research in Huang lab is supported by the W. M. Keck Foundation, the NIH Single Cell Analysis Program (R33EB019784 to B. C. and B.H.) and the NIH Extracellular RNA Communication Consortium (U19CA179512, TO S.B. and B.H.). B.H. is a Chan Zuckerberg Biohub investigator.

